# Assessment of the reproducibility and inter-site transferability of the murine direct splenocyte mycobacterial growth inhibition assay (MGIA)

**DOI:** 10.1101/2021.02.14.431105

**Authors:** Rachel Tanner, Andrea Zelmer, Hannah Painter, Elena Stylianou, Nawamin Pinpathomrat, Rachel Harrington-Kandt, Lucia Biffar, Michael J. Brennan, Helen McShane, Helen A. Fletcher

**Author notes:** Corresponding author, The Jenner Institute, Old Road Campus Research Building, Roosevelt Drive, Headington, Oxford, OX3 7DQ. indicate equal contribution (Rachel Tanner, Andrea Zelmer). indicate equal contribution (Helen McShane, Helen A. Fletcher).

## Abstract

Tuberculosis (TB) vaccine candidates must be tested for safety and efficacy using preclinical challenge models prior to advancement to human trials, because of the lack of a validated immune correlate or biomarker of protection. New, unbiased tools are urgently needed to expedite the selection of vaccine candidates at an early stage of development and reduce the number of animals experimentally infected with virulent *Mycobacterium tuberculosis* (*M*.*tb*). In recent years, there has been a concerted effort to develop standardised functional *ex vivo* mycobacterial growth inhibition assays (MGIAs) as a potential surrogate read-out of vaccine efficacy. We have previously described a direct MGIA for use with mouse splenocytes. In the current study, we set out to systematically compare co-culture conditions for the murine direct splenocyte MGIA with respect to both intra-assay repeatability and inter-site reproducibility. Common sample sets were shared between laboratory sites and reproducibility and sensitivity to detect a BCG-vaccine induced response were assessed. Co-culturing 5×10^6^ splenocytes in 48-well plates resulted in improved reproducibility and superior sensitivity to detect a vaccine response compared with standing or rotating sealed 2ml screw-cap tubes. As the difference between naïve and BCG vaccinated mice was not consistently detected across both sample sets at both sites, we sought to further improve assay sensitivity by altering the multiplicity of infection (MOI). Cell viability at the end of the co-culture period was improved when splenocyte input number was reduced, with the highest viability for the condition of 3×10^6^ splenocytes in 48-well plates. This cell input was also associated with the greatest sensitivity to detect a BCG vaccine-mediated MGIA response using an *M*.*tb* inoculum. Based on our findings, we recommend optimal co-culture conditions in a move towards aligning direct MGIA protocols and generating a cross-species consensus for early evaluation of TB vaccine candidates and biomarker studies.

## 1.0 Introduction

Tuberculosis (TB) remains a major global health problem, with 10 million new cases and 1.4 million deaths in 2019 (WHO, 2020). Bacillus Calmette Guérin (BCG) is the only currently available vaccine against TB. However, BCG confers incomplete and variable protection against pulmonary TB in adolescents and adults, and a new more efficacious TB vaccine is urgently needed (Fine, 1995; Mangtani et al., 2014). Due to the lack of a validated immune correlate or biomarker of protection from TB, candidate vaccines are currently tested for safety and efficacy in long and costly preclinical challenge experiments prior to progression into human trials (McShane and Williams, 2014). New, unbiased tools are urgently needed to refine and expedite the early selection of vaccine candidates. One such potential tool is an *ex vivo* mycobacterial growth inhibition assay (MGIA), which measures a functional ‘sum-of-the-parts’ vaccine response as a potential surrogate of vaccine-induced protection.

A successful MGIA would reduce the number of candidate vaccines going forward to virulent *M*.*tb* challenge experiments, and vaccine efficacy against different clinical isolates could be evaluated using cells from a single group of vaccinated animals, further reducing the number of animals used. Such an assay could also be applied in the measurement of vaccine potency, lot-to-lot consistency and stability as an alternative to *in vivo* experiments. Preclinical MGIAs provide an opportunity for biological assay validation against measures of protection from *in vivo M*.*tb* challenge, which is not ethically possible in humans, and can then be bridged to use with human cells with greater confidence. These goals are in line with the 3Rs principles of reduction, refinement and replacement of the use of animals in scientific research (Burden et al., 2015; Tanner and McShane, 2017).

A number of MGIAs have been described in the literature for use with samples from humans, cattle and mice (reviewed in (Tanner et al., 2016)). However, such assays are technically demanding and often complex, and there is a lack of systematic assessment for reproducibility or technical/biological validation which has limited their uptake within the TB vaccine field. We have worked to develop and qualify a transferable cross-species MGIA (Brennan et al., 2017). Originally adapted from the methods of Wallis et al. (Cheon et al., 2002) and now referred to as the ‘direct MGIA’, the advantages of the assay include its relative simplicity (as PBMC or splenocytes are inoculated with mycobacteria directly rather than culturing macrophages and adding in expanded effector cells), thus improving reproducibility and transferability, and reducing bias towards any particular immune parameter/s. It also makes use of routine reagents and equipment, and the standardised and rapid BACTEC MGIT quantification system rather than traditional colony counting (Brennan et al., 2017).

We have previously optimised, standardised and harmonised the direct MGIA for use with PBMC from humans and non-human primates (NHPs), and we and others have applied it to demonstrate a BCG vaccine-induced improvement in mycobacterial control, and confirm a biologically relevant response that correlates with protection from experimental *in vivo* mycobacterial infection (Fletcher et al., 2013; Smith et al., 2016; Joosten et al., 2018; Prabowo et al., 2019; Tanner et al., 2019; Tanner et al., 2020; Tanner et al., 2021). In parallel, the murine direct MGIA has been applied to detect a response to BCG and other TB vaccine candidates (Marsay et al., 2013; Yang et al., 2016; Zelmer et al., 2016; Jensen et al., 2017) and a preliminary protocol made available on BioRxiv (Zelmer et al., 2015). Importantly, a preclinical MGIA provides the opportunity for biological validation against direct measures of *in vivo* protection from *M*.*tb* challenge, which is not ethically possible in humans. It also permits the testing of efficacy against multiple *M*.*tb* isolates and the role of different immune parameters using a single set of cells, thus reducing the number of animals required (Tanner and McShane, 2017).

We, and others, have described steps towards optimisation and further development of the murine direct MGIA, and several methodological variations have been reported in the literature (Yang et al., 2016; Zelmer et al., 2016; Jensen et al., 2017). Transferring and harmonising a single standardised protocol is critical to ensure that comparable information can be extracted from ongoing and future preclinical vaccine trials conducted across different organisations. The aim of the current study was to systematically compare co-culture conditions for the murine direct splenocyte MGIA across sites and to further develop assay sensitivity in an attempt to achieve an optimised consensus for transfer to different preclinical TB vaccine laboratories.

## 2.0 Methods

### 2.1 In vivo mouse experiments

#### 2.1.1 Mice

Female C57BL/6 or BALB/C (as specified) mice aged 5-8 weeks were obtained from Charles River UK Ltd. (Kent, UK) or Envigo (Cambridgeshire, UK) and acclimatised for a minimum of 5 days at the research facility prior to the first experimental procedure. Experiments were performed at the University of Oxford and the London School of Hygiene and Tropical Medicine (LSHTM) under appropriate UK Home Office licences and in accordance with the UK Home Office Regulations (Guidelines on the Operation of Animals, Scientific Procedures Act, 1986). At Oxford, animals studies were conducted with ethical approval from the local Ethical Review Committee, at LSHTM they were approved by the London School of Hygiene and Tropical Medicine Animal Welfare and Ethics Review Board. Mice were housed in ventilated cages with enrichment and a 12-hour light/dark cycle and controlled temperature (20-22°C), and provided with normal chow and water *ad libitum*.

#### 2.1.2 BCG vaccination and splenocyte harvesting

BCG Pasteur was thawed at room temperature and diluted to a final concentration of 2-5×10^6^ CFU/ml in physiological saline solution (Baxter Healthcare, Newbury, UK). In the route of vaccination experiment (Figure 1), half of the mice (n=8, BALB/c) were randomised to receive 100µl BCG vaccination by the subcutaneous (SC) route into the base of the tail and half (n=8, BALB/c) to receive 100µl BCG vaccination by the intradermal (ID) route into the ear. In each inter-site comparison experiment (Figure 3), half of the mice (n=8, C57BL/6) were randomised to receive 100µl of BCG SC and half (n=8, C57BL/6) to receive physiological saline solution SC into the leg flap. In the final comparison of MGIA splenocyte input (Figure 4), half of the mice (n=6, C57BL/6) were randomised to receive 100µl of BCG SC and half (n=6, C57BL/6) to receive physiological saline solution SC into the leg flap. Animals were rested for 6 weeks, after which time they were euthanised and spleens dissected aseptically. Splenocytes were isolated by mechanical disruption through a 100µm cell strainer. Red blood cells were lysed, and then splenocytes were counted and resuspended in antibiotic-free media (containing 10% foetal calf serum, 2mM l-glutamine and 25mM HEPES) for the direct MGIA.

**Figure 1.**
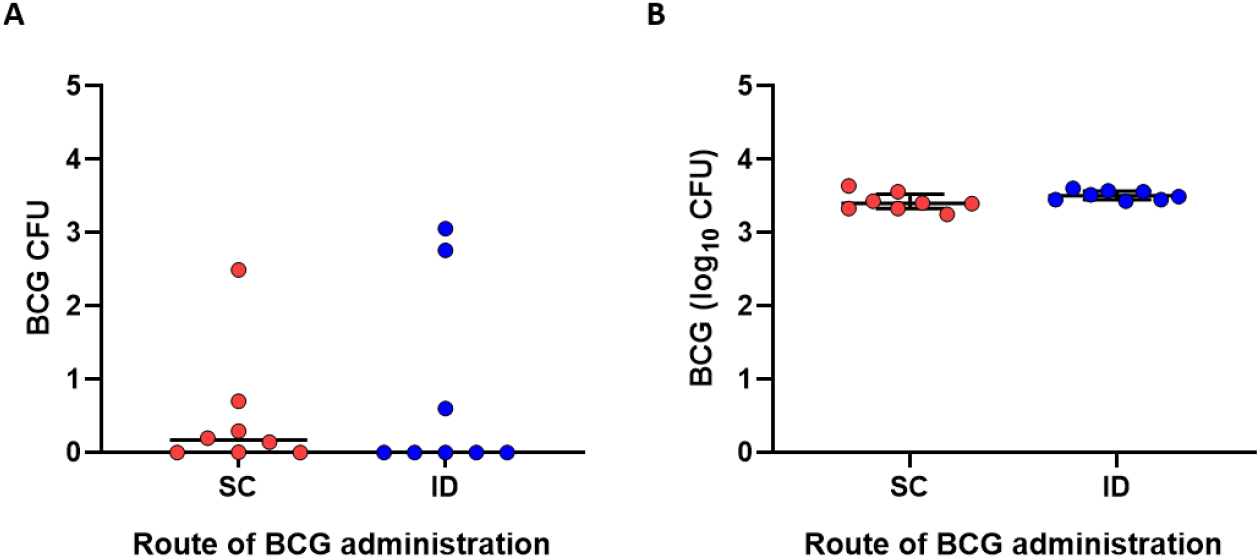
Comparison of route of BCG administration. BALB/c mice were vaccinated with BCG via the subcutaneous (SC, n=8, red) or intradermal (ID, n=8, blue) route. Residual BCG in 5×10^6^ splenocytes removed 6 weeks later was quantified using the BACTEC MGIT system (A), and the direct splenocyte MGIA was performed using 5×10^6^ cells inoculated with ∼100 CFU BCG Pasteur and the rotating in-tube method (B). Points represent mice and lines represent median values with the inter-quartile range (IQR).

**Figure 2.**
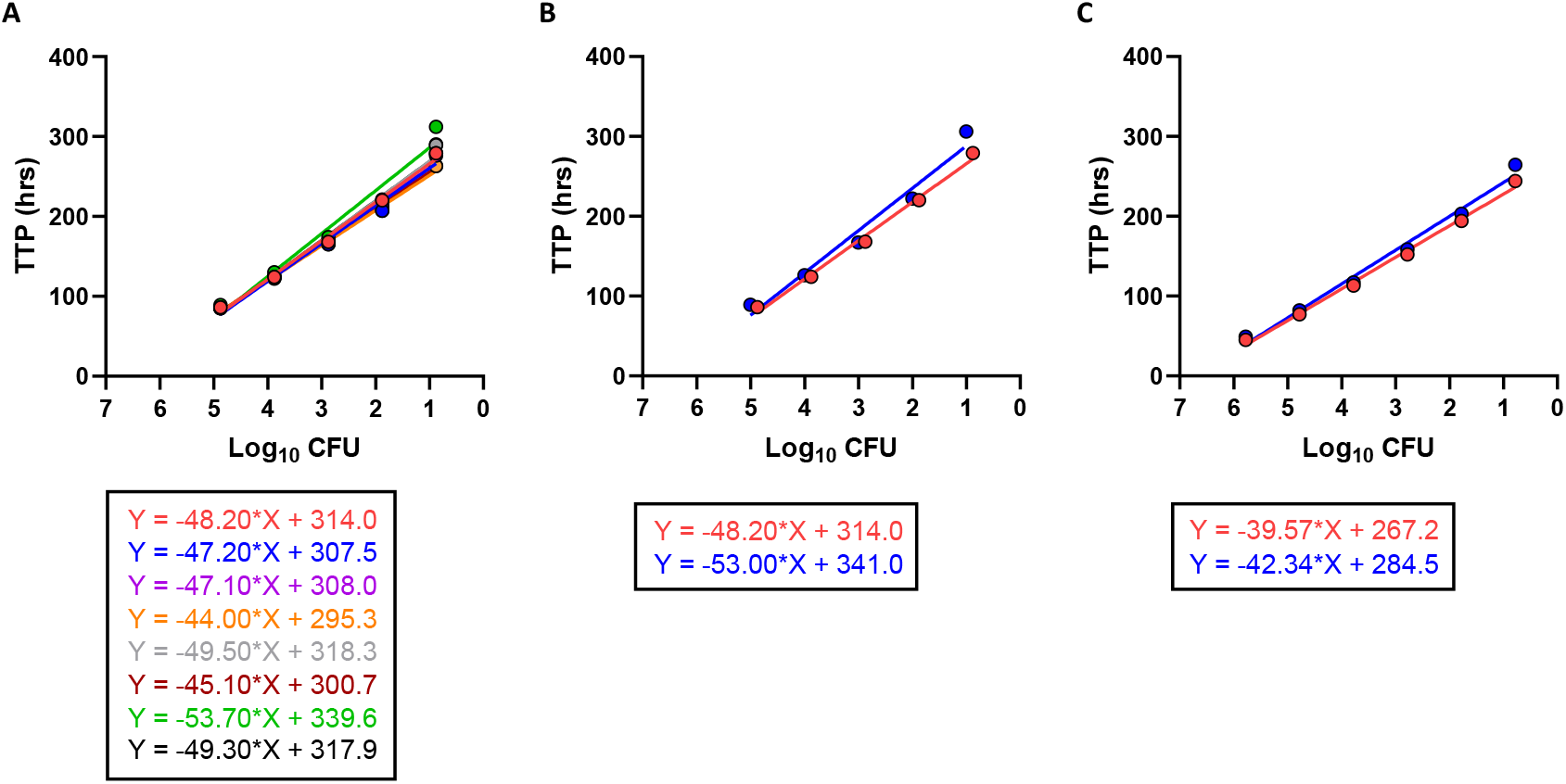
Assessment of standard curve reproducibility. Replicate standard curves for BCG Pasteur Aeras were compared when performed at the same site at the same time (A), at the same site 3 months apart (B), and at two different sites (C). Points represent the mean of duplicate dilution tubes. Linear regression analysis was carried out by fitting a semi-log line and the equations of the curves shown.

**Figure 3.**
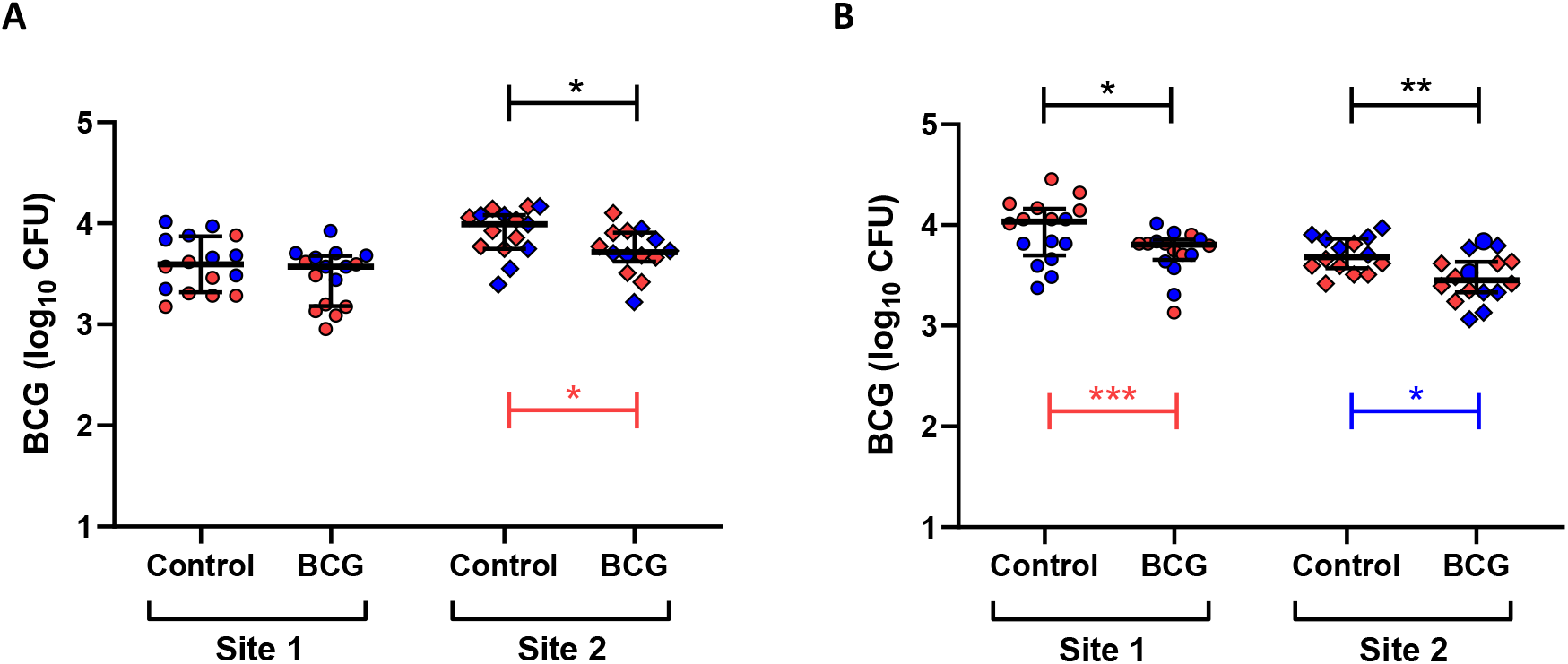
Inter-site comparison of direct splenocyte MGIA methods. The direct splenocyte MGIA was performed on two independent shared sample sets from n=8 C57BL/6 mice per group using either the rotating in-tube protocol (A) or the 48-well plate protocol (B). MGIA outcomes are shown for Site 1 (circles) and Site 2 (diamonds), and for experiment 1 (red) and experiment 2 (blue). Significance is shown for each individual experiment in the corresponding colour, and for both sample sets combined in black. Unpaired t-tests were performed between naïve control and BCG-vaccinated mice for each method at each site, where * indicates a p-value of <0.05, ** indicates a p-value of <0.01, and *** indicates a p-value of <0.001. Points represent individual mice and lines represent median values with the inter-quartile range (IQR).

**Figure 4.**
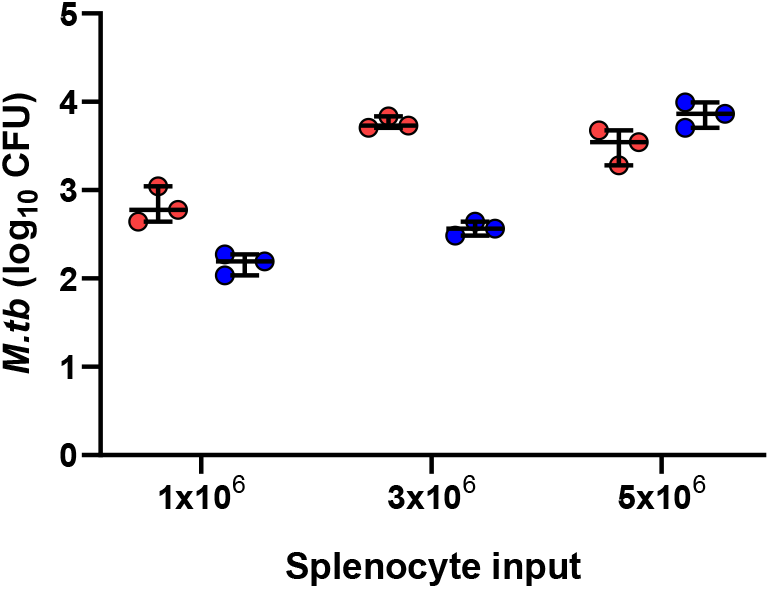
Comparison of splenocyte input. The direct splenocyte MGIA using the 48-well plate method was performed using splenocytes from naïve (red) and BCG vaccinated (blue) C57BL/6 mice at different input numbers inoculated with ∼100 CFU *M*.*tb* Erdman. Points represent individual co-cultures comprising pooled cells from n=6 mice per group and lines represent the median value with the inter-quartile range (IQR).

### 2.2 Direct splenocyte mycobacterial growth inhibition assay (MGIA)

#### 2.2.1 Mycobacteria stocks

BCG Pasteur was obtained from Aeras (Rockville MD, USA) as 500μL frozen aliquots. *M*.*tb* Erdman was obtained from BEI Resources (Manassas, VA, USA) as a 1mL frozen glycerol stock and grown to log phase at 150 rpm and 37°C in Middlebrook 7H9 Broth (Yorlab, York, UK) supplemented with 10% OADC (Yorlab), 0.05% Tween-80 (Sigma, Gillingham, UK) and 0.2% glycerol (Sigma). Mycobacterial stocks were stored at −80 °C. All work using *M*.*tb* Erdman was performed in a biosafety containment level 3 laboratory at LSHTM in accordance with guidance from the UK Advisory Committee on Dangerous Pathogens (ACDP).

#### 2.2.2 Preparation of standard curves

Serial 10-fold dilutions of BCG Pasteur Aeras or *M*.*tb* Erdman stock were prepared in supplemented Middlebrook 7h9 with thorough mixing between each dilution. 500µl of each dilution was added to duplicate BACTEC MGIT tubes supplemented with PANTA antibiotics (polymyxin B, amphotericin B, nalidixic acid, trimethoprim and azlocillin) and Oleic Albumin Dextrose Catalase (OADC) enrichment broth (Becton Dickinson, UK), placed on the BACTEC 960 instrument (Becton Dickinson, UK), and incubated at 37°C until the detection of positivity by fluorescence. In parallel, 3 x 20µl from each dilution was spotted onto sections of 7H11 agar plates and colonies counted after 10-14 days. Standard curves were generated by plotting MGIT time-to-positivity (TTP) against the equivalent CFU for each input volume of BCG. Regression analysis was used to obtain the equation for each semi-log curve.

#### 2.2.3 In-tube protocol

2ml screw-cap tubes containing 1×10^6^, 3×10^6^ or 5×10^6^ splenocytes (as specified) and ∼100 CFU BCG Pasteur in a total volume of 600µl RPMI (containing 10% foetal calf serum, 2mM l-glutamine and 25mM HEPES) were incubated for 96 hours at 37°C either on a 360° rotator (VWR International) or standing stationary (as specified). The co-cultures were then added directly to BACTEC MGIT tubes supplemented with PANTA antibiotics and OADC enrichment broth (Becton Dickinson, UK).

#### 2.2.4 48-well plate protocol

48-well tissue culture plates containing 1×10^6^, 3×10^6^ or 5×10^6^ splenocytes (as specified) and ∼100 CFU BCG Pasteur or *M*.*tb* Erdman (as specified) in a total volume of 600µl RPMI (containing 10% heat-inactivated foetal calf serum, 2mM l-glutamine and 25mM HEPES) per well were incubated for 96 hours at 37°C with CO_2_. At the end of the culture period, co-cultures were added to 2ml screw-cap tubes and centrifuged at 12,000rpm for 10 minutes. During this time, 500µl sterile water was added to each well to lyse adherent monocytes. Supernatants were removed from the 2ml screw-cap tubes, and water from the corresponding well added to the pellet. Tubes were pulse vortexed and the full volume of lysate transferred to BACTEC MGIT tubes supplemented with PANTA antibiotics and OADC enrichment broth (Becton Dickinson, UK). For the final experiment, supernatants from the co-cultures were carefully removed and discarded. 500µl sterile water was added to each well to lyse adherent monocytes, and incubated at room temperature for 5 minutes before transferring the total volume to BACTEC MGIT tubes supplemented with PANTA antibiotics and OADC enrichment broth (Becton Dickinson, UK).

#### 2.2.5 Mycobacterial quantification

At the end of all MGIA protocols, MGIT tubes were placed on the BACTEC 960 instrument (Becton Dickinson, UK) and incubated at 37°C until the detection of positivity by fluorescence. On day 0, duplicate direct-to-MGIT control tubes were set up by inoculating supplemented BACTEC MGIT tubes with the same volume of mycobacteria as the samples. The time to positivity (TTP) read-out was converted to log_10_ CFU using stock standard curves generated as described in 2.2.1.

### 2.3 Statistical analysis

As MGIA data were log transformed and normally distributed, unpaired t-tests were used to compare between different treatment groups. For non-parametric data (for example residual BCG recovery from splenocytes), a Mann Whitney test was used. Intra-class correlation coefficients (two-way mixed model, consistency agreement, single measures) were calculated using raw TTP values to assess intra-assay repeatability between duplicate co-cultures and inter-site reproducibility between shared sample sets assayed at two different institutes. ICC categories for interpreting kappa values were taken from the guidelines of Landis and Koch (Landis and Koch, 1977).

## 3.0 Results

### 3.1 Low levels of residual BCG are recovered from splenocytes following BCG vaccination, but route of administration does not affect outcome of the direct splenocyte MGIA

To determine the most appropriate route of BCG administration for our inter-site experiments, we first assessed MGIA outcome and recovery of residual BCG in splenocytes from mice vaccinated with BCG by either the subcutaneous (SC, n=8) or intradermal (ID, n=8) routes. Residual BCG was recovered in 5 out of 8 mice in the SC group, and 3 out of 8 mice in the ID group, although in all cases the number of CFU were extremely low (approximately 3 logs lower than the number of CFU recovered at the end of the 96 hour MGIA co-culture) in all cases. There was no statistically significant difference between the groups (p=0.4, Mann-Whitney, Figure 1A). There was no difference in control of mycobacterial growth following SC compared with ID vaccination (Δlog_10_ CFU = 0.09; p=0.09, unpaired t-test; Figure 1B).

### 3.2 Assessment of assay reproducibility

To determine assay robustness, reproducibility of the standard curve for BCG Pasteur Aeras stock was evaluated by adding 10-fold serial dilutions to duplicate BACTEC MGIT tubes and in parallel plating on 7H11 agar to generate corresponding time-to-positivity (TTP) and CFU values. Replicate curves demonstrated good consistency when performed eight times at the same site at the same time (Figure 2A), twice at the same site 3 months apart (Figure 2B), or at two different sites (Figure 2C).

Intra-assay repeatability was compared between replicate MGIA co-cultures of 5×10^6^ splenocytes from n=16 C57BL/6 mice inoculated with ∼100 CFU BCG Pasteur across different assay conditions. These included co-culturing in sealed 2ml screw-cap tubes that were either rotating 360° or standing stationary at 37°C for 96 hours (Marsay et al., 2013; Zelmer et al., 2016; Jensen et al., 2017), or co-culturing in 48-well tissue culture plates with CO_2_ (Yang et al., 2016; Painter et al., 2020). In all cases, mycobacteria was quantified at the end of the co-culture period using the BACTEC MGIT liquid culture system. We observed an intraclass correlation coefficient (ICC) value of 0.09 (slight agreement) between replicate co-cultures for standing in-tube, 0.55 (moderate agreement) for rotating in-tube, and 0.77 (substantial agreement) for 48-well plate protocols. Spleens from n=16 mice were halved and shared between two different laboratories for assessment of inter-site reproducibility. ICC values were 0.19 (slight agreement) for standing in-tube, 0.51 (moderate agreement) for rotating in-tube, and 0.42 (moderate agreement) for 48-well plate protocols; data are summarised in Table 1.

**Table 1.**
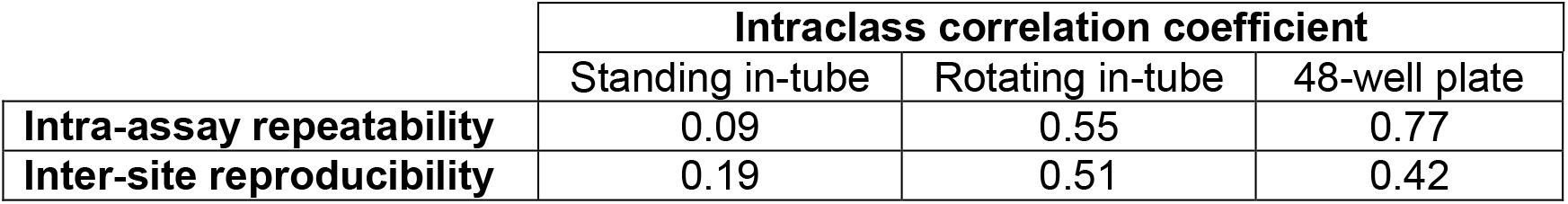
Intraclass correlation coefficients (ICC) for intra-assay repeatability and inter-site reproducibility for direct splenocyte MGIA methods.

### 3.3 Sensitivity to detect a BCG-vaccine induced response across sites is improved by co-culturing in 48-well plates compared with screw-cap tubes

Spleens from n=8 naïve and n=8 BCG vaccinated C57BL/6 mice originating from Site 1 were halved and processed at two different laboratory sites, where the direct splenocyte MGIA was performed independently using 5×10^6^ cells and ∼100 CFU BCG Pasteur per co-culture. In the first experiment, the difference in MGIA outcome was compared between naïve and BCG vaccinated mice using either the rotating in-tube or 48-well plate protocols. Using the rotating in-tube protocol, there was significantly improved control of mycobacterial growth in the BCG vaccinated compared with naïve mice at Site 2 (Δlog_10_ CFU = 0.21; p=0.04, unpaired t-test; Figure 3A). Using the 48-well plate protocol, there was significantly improved control of mycobacterial growth in the BCG vaccinated group at Site 1 (Δlog_10_ CFU = 0.46; p=0.0004, unpaired t-test; Figure 3B).

In the second experiment, the difference in MGIA outcome was compared between naïve and BCG vaccinated mice using either the standing in-tube, rotating in-tube or 48-well plate protocols. Using the standing in-tube protocol, there was no difference in growth control between naïve and BCG vaccinated mice at either site, although there was a trend towards improved control in the BCG vaccinated group at Site 1 (p=0.07, unpaired t-test, data not shown). Using the rotating in-tube protocol, there was no difference at either site (Figure 3A). Using the 48-well plate protocol, there was significantly improved control of mycobacterial growth in the BCG vaccinated group at Site 2 and trend towards the same effect at Site 1 (Δlog_10_ CFU = 0.38 and Δlog_10_ CFU = 0.12; p=0.01 and p=0.09 respectively, unpaired t-test; Figure 3B). When outcomes were combined for the two sample sets, there was significantly improved control of mycobacterial growth in the BCG vaccinated compared with naïve mice at Site 2 using the rotating in-tube protocol (Δlog_10_ CFU = 0.20; p=0.03, unpaired t-test; Figure 3A), and at both sites using the 48-well plate protocol (Δlog_10_ CFU = 0.22, p=0.03 at Site 1 and Δlog_10_ CFU = 0.23, p=0.005 at Site 2, unpaired t-test; Figure 3B).

### 3.4 A reduction in splenocyte number results in improved cell viability and sensitivity to detect a BCG-vaccine induced response

While the 48-well plate method demonstrated superior sensitivity to detect a BCG-vaccine induced response compared with in-tube methods, the difference between naïve and BCG vaccinated mice was not consistently detectable across both experiments at both sites. We therefore sought to further optimise the assay by assessing cell viability at the end of the co-culture period across different splenocyte inputs (n=3 co-cultures for each condition). Viability was greatest using the 48-well plate method for all cell inputs, with the best outcome (median 54%) for an input of 3×10^6^ splenocytes; data are summarised in Table 2.

**Table 2.**
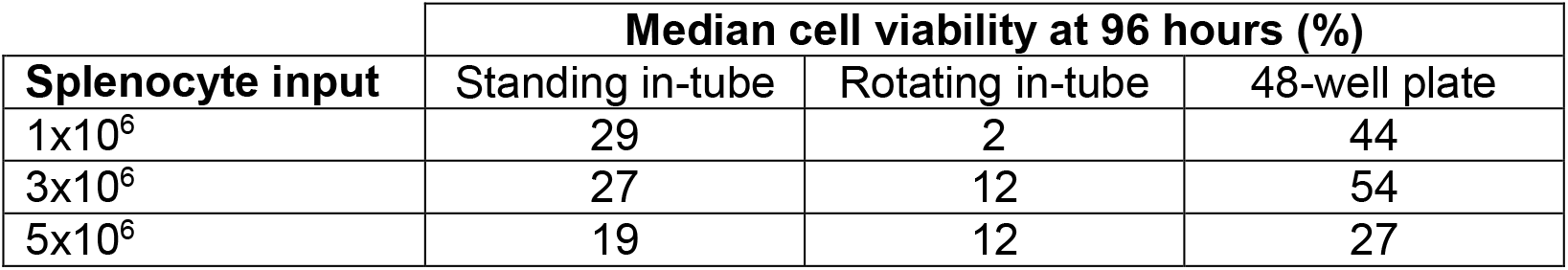
Median cell viability at 96 hours for direct splenocyte MGIA methods.

| Splenocyte input | Median cell viability at 96 hours (%) |  |  |
| --- | --- | --- | --- |
|  | Standing in-tube | Rotating in-tube | 48-well plate |
| $1 \times 10^6$ | 29 | 2 | 44 |
| $3 \times 10^6$ | 27 | 12 | 54 |
| $5 \times 10^6$ | 19 | 12 | 27 |

In a separate experiment of n=16 C57BL/6 mice, intra-assay repeatability between duplicate MGIA co-cultures was assessed for different co-culture conditions (cell number permitting). Using the rotating in-tube assay, the most consistent outcomes were achieved with an input of 3×10^6^ splenocytes (ICC of 0.70, substantial agreement). The best overall outcome by this measure was again the 48-well plate method with 3×10^6^ splenocytes (ICC of 0.91, almost perfect agreement); results are summarised in Table 3. The direct splenocyte MGIA under these conditions showed a significant improvement in control of mycobacterial growth following SC BCG vaccination as previously reported (p=0.001, unpaired t-test) (Painter et al., 2020).

**Table 3.**
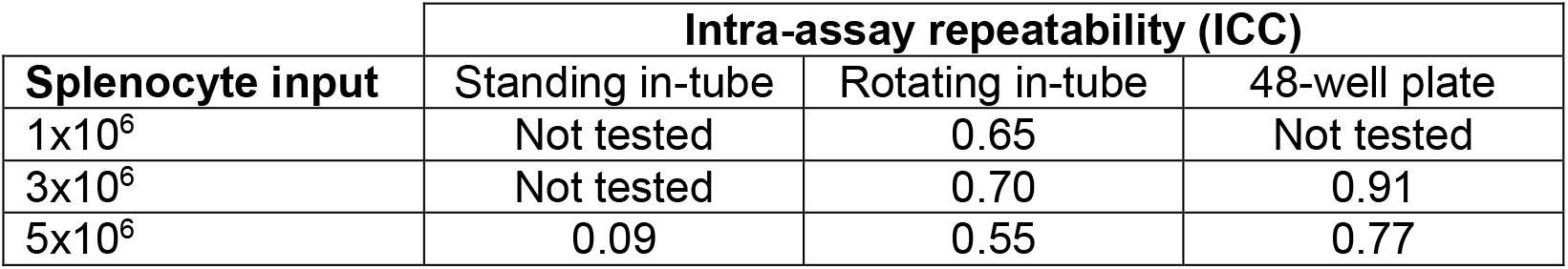
Intra-assay repeatability for direct splenocyte MGIA methods.

Yang et al. and Painter et al. have previously described a method whereby co-cultures are prepared by pooling splenocytes from all mice within a group to improve assay reproducibility and thus sensitivity (Yang et al., 2016; Painter et al., 2020). We therefore compared the variability between 10 co-cultures containing 3×10^6^ splenocytes from either 10 individual mice, or from a pool of splenocytes from the same 10 mice. The mean log_10_ CFU was comparable between conditions (pooled = 3.40 log_10_ CFU, individual = 3.38 log_10_ CFU), but reproducibility was improved for the pooled condition (CV = 3.7%, STDEV = 0.13 log_10_ CFU, range = 0.43 log_10_ CFU) compared with individual mice (CV = 7.6%, STDEV = 0.27 log_10_ CFU, range = 0.91 log_10_ CFU). Using the 48-well plate method and ∼100 CFU *M*.*tb* Erdman as the mycobacterial inoculum, we then compared ability to detect a BCG-vaccine induced response with different splenocyte inputs and 3 replicate co-cultures comprising combined splenocytes from 6 mice per group. There was an increase in mycobacterial growth with increased cell number, and improved control of mycobacterial growth was detected in the BCG vaccinated compared with control mice for splenocyte inputs of 1×10^6^ and 3×10^6^ (Figure 4). The greatest delta between groups was seen for a splenocyte input of 3×10^6^ (Δlog_10_ CFU = 1.19), which was approximately double that for 1×10^6^ cells (Δlog_10_ CFU = 0.65).

## 4.0 Discussion

A validated, reliable and transferable consensus murine MGIA would be an invaluable tool to refine and expedite the evaluation and down-selection of experimental TB vaccine candidates at an early stage of development. Several methodological variations of the direct splenocyte MGIA have been reported in the literature (Yang et al., 2016; Zelmer et al., 2016; Jensen et al., 2017). We sought to systematically compare co-culture conditions for the murine direct splenocyte MGIA across sites and to further develop assay sensitivity in an attempt to achieve a consensus for transfer to different preclinical TB vaccine laboratories.

BCG vaccination in murine studies is routinely administered by the subcutaneous (SC) route, but in humans is most often given intradermally (ID); route of BCG administration has been shown to influence the quantity and quality of the immune response induced (Dockrell and Smith, 2017). Furthermore, persistent live BCG bacilli have been reported in the spleens of mice up to 30 weeks post-vaccination by both SC and ID routes (Olsen et al., 2004; Kaveh et al., 2014). We therefore compared MGIA outcome following BCG vaccination by the two different routes and quantified residual BCG in splenocytes to determine the most appropriate route for our inter-site experiments. There was no difference in mycobacterial growth control between mice receiving SC and ID vaccination, consistent with human studies showing that percutaneous and intradermal routes confer similar immunogenicity and efficacy in infants (Davids et al., 2006; Hawkridge et al., 2008; Lee et al., 2011; Dockrell and Smith, 2017). Interestingly, residual BCG was recovered in splenocytes from each of 5/8 mice following SC vaccination and 3/8 mice following ID vaccination. While up to 3 log_10_ CFU has been recovered from the spleen at 6 weeks post-SC vaccination (Olsen et al., 2004), we sampled just 5×10^6^ splenocytes per mouse.

The presence of residual BCG could have important implications for the MGIA: 1) Such BCG would also grow in MGIA co-cultures and be indistinguishable from the *in vitro* inoculum during the quantification stage, thus potentially masking growth inhibition and reducing assay sensitivity, and 2) the MGIA may be measuring effector responses that have been primed by live replicating BCG rather than memory responses, making BCG a poor bench-mark against which to develop an assay designed to assess efficacy of non-live vaccines. Painter et al. recently described the use of 2-thiophenecarboxylic acid hydrazide (TCH) in the murine direct lung MGIA to prevent the presence of residual BCG following intranasal vaccinationfrom confounding quantification of the MGIA *M*.*tb* Erdman inoculum (Painter et al., 2020). A similar approach could be applied here, although it should be noted that the numbers of BCG recovered were negligible (more than 3 log lower than that quantified at the end of the co-culture period and ∼2 log lower than residual BCG following intranasal vaccination), and therefore unlikely to interfere with the assay, particularly at the lower cell inputs recommended.

Preliminary reproducibility experiments focussed on standard curves generated at the same site at the same time, at the same site 3 months apart, or at two different sites. There was a high level of consistency in curve equations for all comparisons, offering confidence in performance of the BCG stock across sites and over time. Assay reproducibility and sensitivity was then assessed for different direct splenocyte MGIA methods by conducting inter-site experiments with two shared sample sets. Co-culturing in 48-well plates, as reported by Yang et al. (Yang et al., 2016), resulted in superior intra-assay and inter-site reproducibility and sensitivity to detect a BCG-vaccine induced response compared with previously-described in-tube methods. This is consistent with our findings in the human direct PBMC MGIA which demonstrated improved cell viability, IFN-γ production and reproducibility using 48-well plates (Tanner et al., 2019). We also found a similar effect in NHPs (Tanner et al., 2021).

Interestingly, splenocyte viability at the end of the 96 hour co-culture was lower compared with previously-reported values for cryopreserved human PBMC using both the rotating in-tube (27% compared with 50%) and the 48-well plate (54% compared with 73%) methods (Tanner et al., 2019). Murine splenocytes are widely considered more fragile than PBMC, necessitating the use of fresh samples and suggesting a requirement for further optimisation of splenocyte culture conditions. Our observations of poor cell viability using in-tube assay conditions are consistent with those of Jensen et al., who reported 21% viability with rotating co-cultures of 5×10^6^ cells which was increased to 46% by changing to standing cultures and enriching the culture medium (Jensen et al., 2017). Interestingly, we observed only a marginal improvement in viability by removing rotation alone (12 to 19%), suggesting that enrichment of the culture medium with sodium pyruvate, non-essential amino acids and 2-mercaptoethanol may have largely mediated the previously-observed effect. Applying this media enrichment to our optimal conditions in 48-well plates may further improve the 54% viability achieved.

We previously compared different splenocyte input numbers for the rotating in-tube method and found that 5×10^6^ cells resulted in a greater BCG vaccine effect size than 3×10^6^ cells (Zelmer et al., 2016). However, the improved cell viability we observed here using the 48-well plate method may have resulted in loss of sensitivity to consistently detect a vaccine response in our transfer experiment. Indeed, visual inspection of co-cultures revealed yellowing of the culture medium with 5×10^6^ cells, suggesting that the presence of CO_2_ allows more cell growth and replication which exhausts the nutrients available before the end of the 96 hour period if the cell input is high. Power to detect a difference between naïve and BCG vaccinated mice using 5×10^6^ cells in 48-well plates was increased when results from the two laboratories were combined to give a group size of n=16 mice, suggesting that the relatively low sensitivity of the assay may necessitate larger group sizes; the 3Rs implications of this should be balanced against advantages of avoiding *in vivo M*.*tb* challenge. The improved reproducibility achieved by pooling cells from mice within a group may provide an alternative route to increased power. Cell viability at the end of the co-culture was considerably greater with 3×10^6^ cells compared with 5×10^6^ cells (54 vs. 27%), and this input was associated with the greatest effect size between naïve and BCG vaccinated mice when tested using an *M*.*tb* inoculum of ∼100 CFU. A similar MOI comparison in 48-well plates using a BCG inoculum is now required.

In conclusion, we addressed our initial aims of assessing reproducibility and inter-site transferability of the murine direct splenocyte MGIA, and attempted to further optimise assay conditions. Our findings suggest favourable direct splenocyte co-culture conditions of 3×10^6^ cells inoculated with ∼100 CFU *M*.*tb* in 48-well plates. Using 3×10^6^ cells in 48-well plates inoculated with ∼500 CFU of BCG, we recently observed an association between direct MGIA outcome and measures of protection from *in vivo* mycobacterial infection in both humans (Tanner et al., 2019) and macaques (Tanner et al., 2021), thus moving towards aligned direct MGIA protocols for a cross-species consensus. The advantages of co-culturing in 48-well plates are biological (presence of CO_2_, increased cell viability) as well as practical (improved feasibility as it negates the requirement for a 360° tube rotator in an incubator). Pooling splenocytes to reduce variability improves power, thus reducing the number of co-cultures and MGIT tubes required, and therefore associated time and costs. Future work should aim to assess inter-site transferability of the final assay conditions. We are now prepared to apply the murine direct splenocyte MGIA in assessing biological validity against preclinical and clinical protection results, comparing live and non-live vaccines and among laboratories and systems.

## 5.0 Acknowledgements

We are grateful to the staff at the animal facilities and to Aeras for providing and distributing the standardised BCG Pasteur stock used in this project. We would also like to thank Henry Dunne for his technical assistance, and Yashica Gupta and Mercy Mugo (In2ScienceUK) for assistance with cell viability studies.

## 6.0 Funding

This work was funded by Aeras grant no. MCA004. HMcS is a Wellcome Trust Senior Clinical Research Fellow and a Jenner Institute Investigator.

## 7.0 Declaration of interest

Declarations of interest: none.

## Notes

### Competing Interest Statement

The authors have declared no competing interest.

## References

Brennan, M.J., Tanner, R., Morris, S., Scriba, T.J., Achkar, J.M., Zelmer, A., Hokey, D.A., Izzo, A., Sharpe, S., Williams, A., Penn-Nicholson, A., Erasmus, M., Stylianou, E., Hoft, D.F., McShane, H. and Fletcher, H.A., 2017, The Cross-Species Mycobacterial Growth Inhibition Assay (MGIA) Project, 2010-2014. Clin Vaccine Immunol 24.

Burden, N., Chapman, K., Sewell, F. and Robinson, V., 2015, Pioneering better science through the 3Rs: an introduction to the national centre for the replacement, refinement, and reduction of animals in research (NC3Rs). Journal of the American Association for Laboratory Animal Science: JAALAS 54, 198–208.

Cheon, S.H., Kampmann, B., Hise, A.G., Phillips, M., Song, H.Y., Landen, K., Li, Q., Larkin, R., Ellner, J.J., Silver, R.F., Hoft, D.F. and Wallis, R.S., 2002, Bactericidal activity in whole blood as a potential surrogate marker of immunity after vaccination against tuberculosis. Clin Diagn Lab Immunol 9, 901–7.

Davids, V., Hanekom, W.A., Mansoor, N., Gamieldien, H., Gelderbloem, S.J., Hawkridge, A., Hussey, G.D., Hughes, E.J., Soler, J., Murray, R.A., Ress, S.R. and Kaplan, G., 2006, The effect of bacille Calmette-Guérin vaccine strain and route of administration on induced immune responses in vaccinated infants. J Infect Dis 193, 531–6.

Dockrell, H.M. and Smith, S.G., 2017, What Have We Learnt about BCG Vaccination in the Last 20 Years? Front Immunol 8, 1134.

Fine, P.E., 1995, Variation in protection by BCG: implications of and for heterologous immunity. Lancet 346, 1339–45.

Fletcher, H.A., Tanner, R., Wallis, R.S., Meyer, J., Manjaly, Z.R., Harris, S., Satti, I., Silver, R.F., Hoft, D., Kampmann, B., Walker, K.B., Dockrell, H.M., Fruth, U., Barker, L., Brennan, M.J. and McShane, H., 2013, Inhibition of mycobacterial growth in vitro following primary but not secondary vaccination with Mycobacterium bovis BCG. Clin Vaccine Immunol 20, 1683–9.

Hawkridge, A., Hatherill, M., Little, F., Goetz, M.A., Barker, L., Mahomed, H., Sadoff, J., Hanekom, W., Geiter, L., Hussey, G. and South African, B.C.G.t.t., 2008, Efficacy of percutaneous versus intradermal BCG in the prevention of tuberculosis in South African infants: randomised trial. BMJ (Clinical research ed.) 337, a2052–a2052.

Jensen, C., Lindebo Holm, L., Svensson, E., Aagaard, C. and Ruhwald, M., 2017, Optimisation of a murine splenocyte mycobacterial growth inhibition assay using virulent Mycobacterium tuberculosis. Scientific Reports 7, 2830.

Joosten, S.A., Meijgaarden, K.E.V., Arend, S.M., Prins, C., Oftung, F., Korsvold, G.E., Kik, S.V., Arts, R.J., Crevel, R.V., Netea, M.G. and Ottenhoff, T.H., 2018, Mycobacterial growth inhibition is associated with trained innate immunity. J Clin Invest.

Kaveh, D.A., Carmen Garcia-Pelayo, M. and Hogarth, P.J., 2014, Persistent BCG bacilli perpetuate CD4 T effector memory and optimal protection against tuberculosis. Vaccine 32, 6911–8.

Landis, J.R. and Koch, G.G., 1977, The measurement of observer agreement for categorical data. Biometrics 33, 159–74.

Lee, H., Cho, S.N., Kim, H.J., Anh, Y.M., Choi, J.E., Kim, C.H., Ock, P.J., Oh, S.H., Kim, D.R., Floyd, S. and Dockrell, H.M., 2011, Evaluation of cell-mediated immune responses to two BCG vaccination regimes in young children in South Korea. Vaccine 29, 6564–71.

Mangtani, P., Abubakar, I., Ariti, C., Beynon, R., Pimpin, L., Fine, P.E., Rodrigues, L.C., Smith, P.G., Lipman, M., Whiting, P.F. and Sterne, J.A., 2014, Protection by BCG vaccine against tuberculosis: a systematic review of randomized controlled trials. Clin Infect Dis 58, 470–80.

Marsay, L., Matsumiya, M., Tanner, R., Poyntz, H., Griffiths, K.L., Stylianou, E., Marsh, P.D., Williams, A., Sharpe, S., Fletcher, H. and McShane, H., 2013, Mycobacterial growth inhibition in murine splenocytes as a surrogate for protection against Mycobacterium tuberculosis (M. tb). Tuberculosis (Edinb) 93, 551–7.

McShane, H. and Williams, A., 2014, A review of preclinical animal models utilised for TB vaccine evaluation in the context of recent human efficacy data. Tuberculosis (Edinb) 94, 105–10.

Olsen, A.W., Brandt, L., Agger, E.M., van Pinxteren, L.A. and Andersen, P., 2004, The influence of remaining live BCG organisms in vaccinated mice on the maintenance of immunity to tuberculosis. Scand J Immunol 60, 273–7.

Painter, H., Prabowo, S.A., Cia, F., Stockdale, L., Tanner, R., Willcocks, S., Reljic, R., Fletcher, H.A. and Zelmer, A., 2020, Adaption of the ex vivo mycobacterial growth inhibition assay for use with murine lung cells. Scientific Reports 10, 3311.

Prabowo, S.A., Smith, S.G., Seifert, K. and Fletcher, H.A., 2019, Impact of individual-level factors on Ex vivo mycobacterial growth inhibition: Associations of immune cell phenotype, cytomegalovirus-specific response and sex with immunity following BCG vaccination in humans. Tuberculosis 119, 101876.

Smith, S.G., Zelmer, A., Blitz, R., Fletcher, H.A. and Dockrell, H.M., 2016, Polyfunctional CD4 T-cells correlate with in vitro mycobacterial growth inhibition following Mycobacterium bovis BCG-vaccination of infants. Vaccine 34, 5298–5305.

Tanner, R. and McShane, H., 2017, Replacing, reducing and refining the use of animals in tuberculosis vaccine research. ALTEX 34, 157–166.

Tanner, R., O’Shea, M.K., Fletcher, H.A. and McShane, H., 2016, In vitro mycobacterial growth inhibition assays: A tool for the assessment of protective immunity and evaluation of tuberculosis vaccine efficacy. Vaccine.

Tanner, R., Satti, I., Harris, S.A., O’Shea, M.K., Cizmeci, D., O’Connor, D., Chomka, A., Matsumiya, M., Wittenberg, R., Minassian, A.M., Meyer, J., Fletcher, H.A. and McShane, H., 2020, Tools for Assessing the Protective Efficacy of TB Vaccines in Humans: in vitro Mycobacterial Growth Inhibition Predicts Outcome of in vivo Mycobacterial Infection. Frontiers in Immunology 10, 2983.

Tanner, R., Smith, S.G., van Meijgaarden, K.E., Giannoni, F., Wilkie, M., Gabriele, L., Palma, C., Dockrell, H.M., Ottenhoff, T.H.M. and McShane, H., 2019, Optimisation, harmonisation and standardisation of the direct mycobacterial growth inhibition assay using cryopreserved human peripheral blood mononuclear cells. J Immunol Methods.

Tanner, R., White, A.D., Boot, C., Sombroek, C.C., O’Shea, M.K., Wright, D., Hoogkamer, E., Bitencourt, J., Harris, S.A., Sarfas, C., Wittenberg, R., Satti, I., Fletcher, H.A., Verreck, F.A.W., Sharpe, S.A. and McShane, H., 2021, A non-human primate in vitro functional assay for the early evaluation of TB vaccine candidates. npj Vaccines 6, 3.

WHO. 2020 World Health Organisation Global tuberculosis report 2020. In

Yang, A.L., Schmidt, T.E., Stibitz, S., Derrick, S.C., Morris, S.L. and Parra, M., 2016, A simplified mycobacterial growth inhibition assay (MGIA) using direct infection of mouse splenocytes and the MGIT system. J Microbiol Methods 131, 7–9.

Zelmer, A., Tanner, R., Stylianou, E., Damelang, T., Morris, S., Izzo, A., Williams, A., Sharpe, S., Pepponi, I., Walker, B., Hokey, D.A., McShane, H., Brennan, M. and Fletcher, H., 2016, A new tool for tuberculosis vaccine screening: Ex vivo Mycobacterial Growth Inhibition Assay indicates BCG-mediated protection in a murine model of tuberculosis. BMC Infect Dis 16, 412.

